# Sum-h^2^, enabling genetic discovery for deep learning-derived phenotypes through a fast evaluation framework and arena of performance

**DOI:** 10.64898/2026.09.24.753959

**Authors:** Tian Xia, Ziqian Xie, Xingzhong Zhao, Sheikh Muhammad Saiful Islam, Ardalan Naseri, Zhiwen Fan, Han Chen, Degui Zhi

**Author notes:** Corresponding author Correspondence to: Dr. Degui Zhi.

## Abstract

In the recent growing interest of AI research toward biology, genetic association studies of AI-derived phenotypes from high-content modalities such as images emerges as a powerful means for biological discovery. However, such AI-phenotyping methods still lacks a good optimization target and an efficient evaluation framework. The number of discovered loci was the major criterion for evaluating the quality of AI-derived phenotypes. However, the time and computational resources required for running the GWAS and subsequent loci-clumping are substantial, limiting the rapid development and iteration of deep learning representation algorithms. Here we present a 1000× faster and lightweight framework, sum-h^2^, than traditional GWAS framework for evaluating genetic discovery through total heritability. We revisit tr(*P*^−1^*G*), previously proposed in the context of evolutionary studies, as a measure of multi-phenotype heritability. We showed that sum-h^2^, the sum of heritability over phenotypic Principal Components (PCs), is equivalent to the linear transformation-invariant tr(*P*^−1^*G*), through both theoretical proof and simulation studies. Moreover, sum-h^2^ can be estimated rapidly with minimal information loss over a relatedness-enriched sample, while preserving the relative ranking of endophenotypes by GWAS loci counts. Based on selected UKB data and sum-h^2^, we set up a Genetic Discovery Arena, enabling rapid and fair comparisons for the development and optimization of deep learning-derived phenotyping methods.

## Introduction

Recently, AI research has increasingly expanded into biology and medicine, as exemplified by efforts from OpenAI and Anthropic to support drug discovery and broader biological research, as well as the emergence of AI agents for automating bioinformatic research [1-3]. However, genetic discovery, a successful approach for unveiling genetic architecture and a key component of early target identification for drug discovery, still lacks standardized and efficient objectives and evaluation frameworks for assessing performance. This limitation hinders the systematic optimization of AI methods for genetic discovery and, consequently, the broader application of AI to this area.

Currently, the most widely used metric for evaluating the genetic discovery potential of newly derived traits is the number of genome-wide significant loci, typically obtained from univariate or multivariate GWAS followed by locus-clumping procedures such as FUMA [4]. This process can require several hours even for a set of 128 traits, limiting its utility for rapid evaluation and optimization. Alternatively, heritability [5-8], which quantifies the proportion of phenotypic variance attributable to genetic variation, can be estimated more efficiently, but is typically evaluated for individual traits. As modern genetic studies increasingly expand toward high-dimensional and multivariate phenotypes, there remains a substantial gap in both (1) a theoretical framework for multivariate heritability that properly accounts for the correlation structure among traits and (2) a practical and scalable method for its estimation.

Here we measure the total heritability 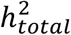 with 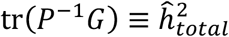for the multivariate traits from the biological evolution field [9]. Here *P* represents the total phenotypic covariance matrix across the k traits/features, where as *G* represents the additive genetic covariance matrix across those same traits/features. Paradoxically, the natural summary is not 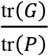, which is sensitive to trait scaling and ignores the orientation of phenotypic correlations: rescaling height from centimeters to meters can radically alter it. Hayes and Hill (1980) [10] recognized this and introduced a *canonical transformation* under which the reparameterized P becomes the identity and G becomes diagonal, with the diagonal elements, the eigenvalues of *Gv* =λ*Pv*, explicitly identified as the heritability of the transformed variables. The quantity tr(*P*^−1^*G*)is the sum of these canonical heritability. It corresponds to the multivariate-heritability branch of the Hansen and Houle (2008) [11] framework, in which the Lande-Arnold equation is standardized by phenotypic variance,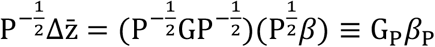(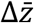represent response to selection while β represent directional selection gradient) after it has been standardized by phenotypic variance), so that G_P_ acts as the multivariate analog of h^2^ and tr(G_P_)=tr(*P*^−1^*G*)summarizes total heritable signal. Ge et al [12] proposed a multidimensional heritability framework for neuroanatomical shape. However, their implementation defines global heritability as the ratio of total variances,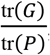, making the metric inherently scale-dependent and sensitive to arbitrary feature scaling. As a result, comparisons of global heritability across different phenotypic representations, feature sets, or learned embeddings can be confounded when those representations differ in scale or variance structure. In contrast, we identify tr(*P*^−1^*G*)as the appropriate measure of total multivariate heritability because it is invariant to nonsingular linear transformations of the phenotype. More recently, Naqvi et al also found that tr(*P*^−1^*G*)is an appropriate multivariate generalization of heritability [13] since it satisfies the following four properties: (1) invariance to units of measurement, (2) coordinate-free, (3) linear in *G*, and (4) maximized with a value of 1 when *G* = *P*, even though they majorly used tr(*P*^−1^*G*)/*k*, where k is the number of multivariate dimensions.

For low-dimensional traits, tr(*P*^−1^*G*)can be calculated directly. However, in high-dimensional settings, the computation becomes increasingly expensive, while the estimation of *P*^−1^ may suffer from singularity or ill-conditioning, introducing additional estimation error. Here, we developed a framework, termed **sum-h**^**2**^, that calculate tr(*P*^−1^*G*)without requiring explicit inversion of *P*. Our method proceeds in two stages: first, we perform a phenotypic PCA on the raw trait matrix to project the correlated phenome onto a set of orthogonal, uncorrelated axes. Second, we estimate the heritability of each resulting PC using Haseman-Elston (HE) regression [14]. HE regression output a lower bound for the narrow-sense heritability h^2^ [15] and is mathematically equivalent to the LD score regression under certain conditions [16, 17]. Our application uses the residual phenotype which is similar to the application within phenotype-correlation genetic-correlation (PCGC) regression [18]. We proved by mathematical derivation and by simulation that the sum of their individual heritability 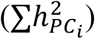 is exactly equivalent to the linear-invariant 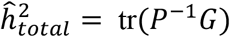. By utilizing HE regression, a method of moments estimator, this framework replaces computationally intensive multivariate genomic-relatedness-based restricted maximum likelihood (GREML) estimation with an efficient, parallelizable workflow that completes in minutes on a standard workstation by exploiting the separability of HE regression across orthogonal PCs.

To our best knowledge, this is the **first** unified benchmarking framework of genetic discovery using a common, representation-invariant metric tr(*P*^−1^*G*), in which phenotype representations can be evaluated 1000x faster than the traditional GWAS framework for total heritable signal. This new metric would allow AI community to rapidly evaluate, iterate, and optimize representation models of the medical images and more, facilitating the development of phenotype representations that better capture biologically meaningful and genetically informative structure. We are establishing a **Genetic Discovery Arena** in which researchers can systematically benchmark their models against a common reference and compare their genetic discovery performance with existing approaches.

## Results

### Theory/Rationale

We define multivariate heritability as tr(*P*^−1^*G*), which is invariant under invertible linear transformations of the phenotype space. When phenotypes are whitened (e.g., PCA with unit variance), this reduces to the sum of per-component heritabilities. Below is the point to point prove.

### a. Why 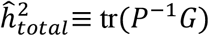 is linear invariant

When we apply a non-singular linear transformation *A* to our traits (i.e., *Y*^′^ =*AY*), the phenotypic (*P*) and genetic (*G*) covariance matrices transform as:

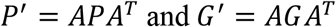

The product *P*^−1^*G* transforms via a **similarity transformation**:

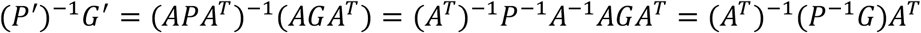

Because *tr*(*ABC*)=*tr*(*BCA*):

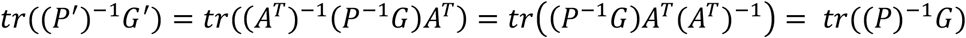

This definition could be further extended to be normalized between 0 and 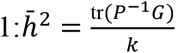, where *k* is the number of traits.

### b. How is tr(***P***^−**1**^***G***)**connect to the sum of *h***^**2**^

When we perform PCA on the phenotype, we are finding the eigenvectors (*e*_*i*_) and eigenvalues (λ_*i*_) of the phenotypic matrix *P*. These eigenvectors define the phenotypic PCs.

- Phenotypic variance of PC *i*: λ_*i*_ (by definition of PCA).
- Genetic variance of PC *i*

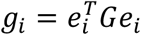

- Heritability of PC *i*

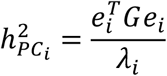

Because the trace of a product of matrices is invariant under cyclic permutations, we can show:

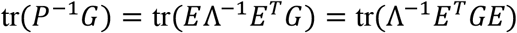

The diagonal elements of the resulting matrix are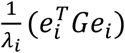, which is exactly 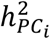. Therefore:

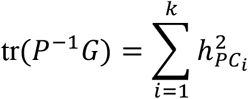

This theoretical derivation is validated by a simulation study (Supplementary Note 1).

### c. How good does 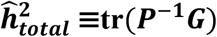 measure the real total heritability 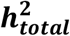 for multivariate phenotypes

From the simulation (Supplementary Note 1), the tr(*P*^−1^*G*)(mean 56.3 across 5 seeds) is close but not equal to the real total heritability (128*0.5=64). Upon further invesigation, we derive a first order coefficient 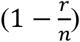 to correct between the total heritability measurement tr(*P*^−1^*G*)and the real total heritability, where the r is rank of 128 and n is the sample size of 2,158. The more accurate correction could be achieved by a fixed-point inversion method. But this coefficient would not affect the rank for different phenotype inputs to the arena, as long as the phenotype dimension and the sample size are the same. The whole simulation process and the correction coefficient deriviation could be found in the Supplementary Note 1.

### d. Why the PCA for 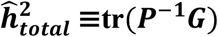is needed

For our original, correlated traits, the heritability of trait *i* is:

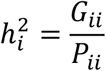

The sum of these is 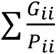. Generally, 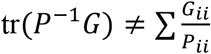 (unless *P* is diagonal)

When *P*^′^ is diagonal:

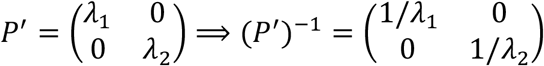

Then, the diagonal elements of (*P*^′^)^−1^*G*^′^ become: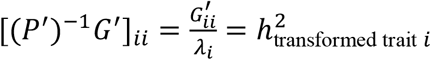. On this specific basis, the trace is exactly the sum of the heritability: 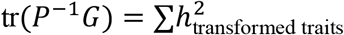

### e. How our framework sum-h^2^ connects with previous work

This sum-h^2^ idea has a direct antecedent in the principal components of heritability (PCH) framework [19], which seeks linear combinations of phenotypes (defined by weight vectors *v*) that maximize heritability, 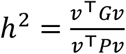. Maximizing this ratio yields the generalized eigenproblem *Gv* =µ*Pv*, where left-multiplying by *v*^⊤^ demonstrates that each eigenvalue µ_*j*_ is exactly the heritability of its corresponding component. Assuming *P* is invertible, we can left-multiply by *P*^−1^ to yield the standard eigenproblem *P*^−1^*Gv* =µ*v*. Thus, the eigenvalues {µ_*j*_} of *P*^−1^*G* are exactly the PCH heritability. Because the trace of any matrix equals the sum of its eigenvalues, it immediately follows that tr(*P*^−1^*G*)=∑_*j*_ µ_*j*_, meaning the total multivariate heritability is simply the sum of these per-axis heritability. While Ott and Rabinowitz explicitly solved this generalized eigenproblem, they viewed it strictly as a dimension-reduction tool to extract the top heritable components for linkage analysis, rather than proposing the sum as an aggregate estimate.

Because the trace is invariant under any change of basis, this total is preserved even when we decompose along phenotypic principal components of variance (PCVs) rather than PCH axes— the per-component heritability differ individually, but their sum is identically tr(*P*^−1^*G*). Further, any full orthogonal decomposition of the phenotypic space yields the identical total.

Our sum-h^2^ approach leverages this by using standard phenotypic PCA followed by per-component Haseman-Elston (HE) regression. This provides a computationally highly efficient route to tr(*P*^−1^*G*)that completely avoids the burden of calculating the full *k* × *k* genetic covariance matrix *G*, while its fundamental equivalence to the PCH decomposition places the estimand on a rigorous and established theoretical footing.

### Framework

We developed a computationally efficient framework to quantify the total heritable signal captured by multidimensional phenotype representations (Figure 1). We first selected a subset of UK Biobank participants enriched for genetic relatedness based on the genomic relationship matrix (GRM; kinship > 0.022) and matched these individuals to their corresponding phenotype representations. Each representation contained up to 128 dimensions (users could use any dimension-reduction method). Phenotypes were demeaned or standardized across individuals and subsequently decorrelated using PCA. HE regression was then performed independently for each PC using GCTA and a precomputed dense GRM, with adjustment for demographic, genetic and imaging-related covariates. The resulting PC-specific heritability were summed across components to obtain a single summary measure, termed **sum-h**^**2**^, representing the aggregate heritable signal captured by the phenotype representation.

**Figure 1.**
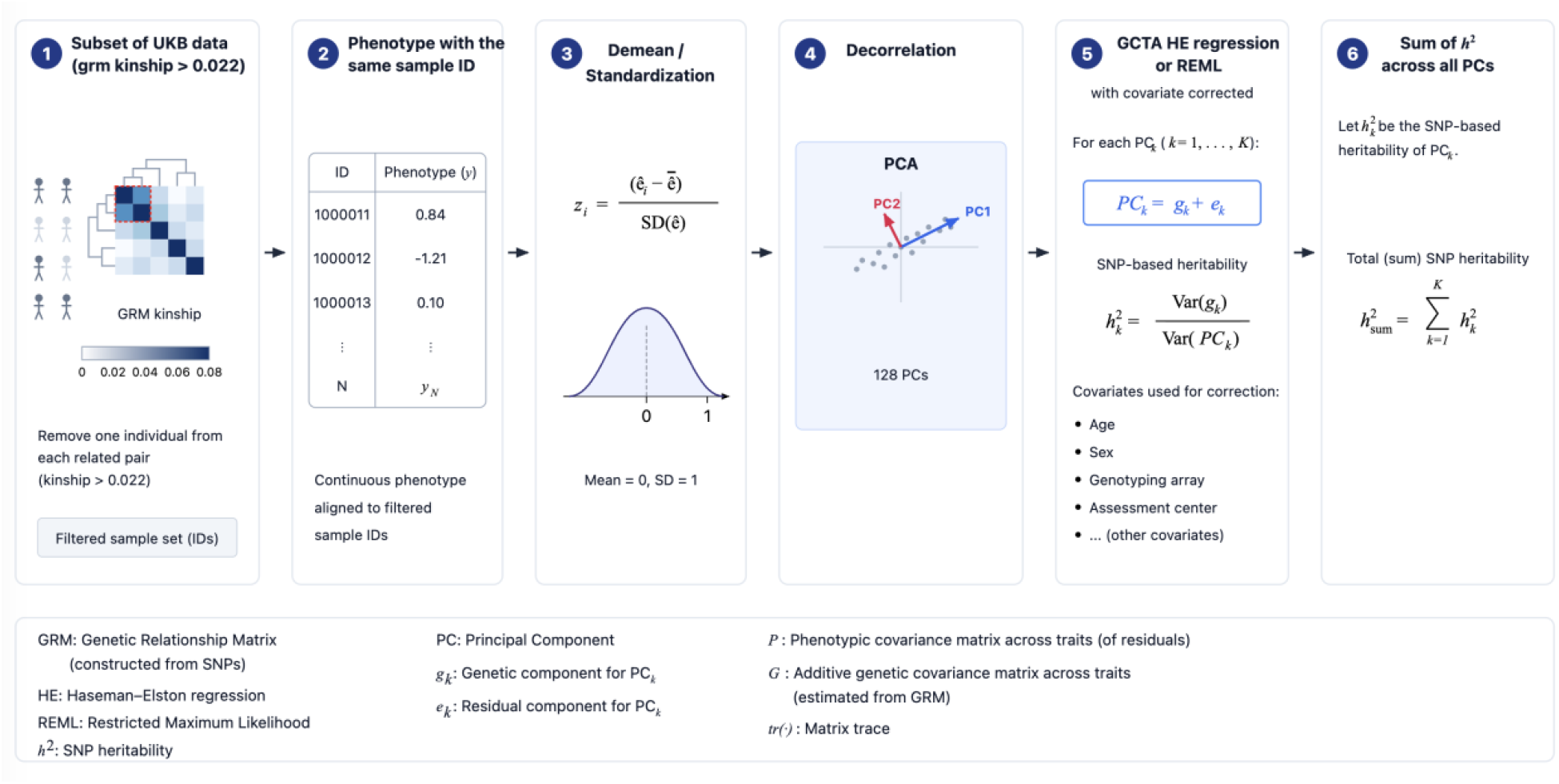
Sum-h^2^framework for deriving total heritability.

The framework requires three primary inputs: a multidimensional phenotype representation with sample identifiers matched to the genetically related UKB subset, a precomputed dense GRM for HE regression, and a corresponding covariate matrix (Table 1). Although we used representations of up to 128 dimensions in the present analysis, the framework can accommodate alternative dimension-reduction strategies before HE regression.

**Table 1.** Required inputs of sum-h^2^ framework.

| Required Inputs | Description |
| --- | --- |
| Phenotype | Phenotype matrix with sample IDs corresponding to the selected UKB subset; dimension $\leq 128$. Users may apply other appropriate dimension-reduction methods. |
| Dense GRM | Precomputed dense genetic relationship matrix (GRM) for the selected UKB subset, used for HE regression. |
| Covariates | Age (field 21003), age<sup>2</sup>, sex (field 31), sex × age, sex × age<sup>2</sup>, 10 genetic PCs (field 22009), head size (field 25000), inverted contrast-to-noise ratio (field 25735), head position in the scanner (fields 25756–25758), scanner table position (field 25759), assessment center location (field 54), and assessment date (field 53). The covariates are the same as those used in our previous publication [20, 21]. |

Covariate adjustment followed our previous study [20, 21] and included age (UKB field 21003), age^2^, sex (field 31), sex-by-age and sex-by-age^2^ interactions, the first ten genetic principal components (field 22009), head size (field 25000), inverted contrast-to-noise ratio (field 25735), scanner head position (fields 25756–25758), scanner table position (field 25759), assessment centre (field 54) and assessment date (field 53). Thus, each phenotype representation was evaluated under an identical genetic background, sample set and covariate-adjustment procedure, enabling direct comparison of the total heritable signal captured by different representations.

We subset the UKB data because HE regression benefits from a dense GRM for more accurate heritability estimation. Selecting samples with GRM kinship > 0.022 enriches off-diagonal GRM values and yields 2,158 samples within the previously used GRM discovery cohort [20, 21]. We selected 128 dimensions as the primary Arena track to reduce computational burden and provide a standardized representation budget for fair comparison. This dimensionality was also supported empirically by our laboratory experiments on deep learning–derived representations of brain MRI, in which 128-dimensional embeddings yielded the largest number of genome-wide significant loci [20, 21].

The framework is highly portable, requiring only a ∼9 MB GRM file and approximately 14 MB for the complete package. We aim to deploy this framework on the UK Biobank RAP platform so that interested groups can evaluate the heritability of their own phenotypes and accommodate the phenotype patient id difference. We also host the results in a public “Arena” on Hugging Face (huggingface.co/spaces/no1summmer/Total_heritiability_ARENA), conceptually similar to large language model leaderboards, with the goal of encouraging improved brain representations for genetic discovery. The complete package is available on GitHub (repository link: https://github.com/ZhiGroup/sum-h2).

### Validation

We evaluated the accuracy of the proposed total heritability metric, calculated by the summed heritability across PCs (sum-h^2^), against the current gold standard measure of genetic discovery, namely the number of identified genomic loci.

More specifically, locus counts were obtained by applying fastGWA analysis to phenotypes from the full UKB discovery cohort [22]. We then used the JAGWAS (with LMM-only features, see methods) [23, 24] approach to aggregate multivariate phenotype summary statistics, followed by FUMA [4] to group significant SNPs into genomic loci. This analysis framework is consistent with our previous publication [20, 21, 24].

To validate the proposed metric, we included all models from the model zoo developed by our group, spanning a wide range of genetic discovery performance (we plan to further validate these findings using external models and loci discovery results, but are currently limited by the availability of compatible endophenotypes with sensitive patient id, which should improve as this approach gains broader adoption). The results have shown that there is a strong correlation between our proposed total heritability metric of sum h^2^ and the number of identified genomic loci with the PCA decorrelation method to 128 dim (Pearson r = 0.925) (Figure 2a).

**Figure 2.**
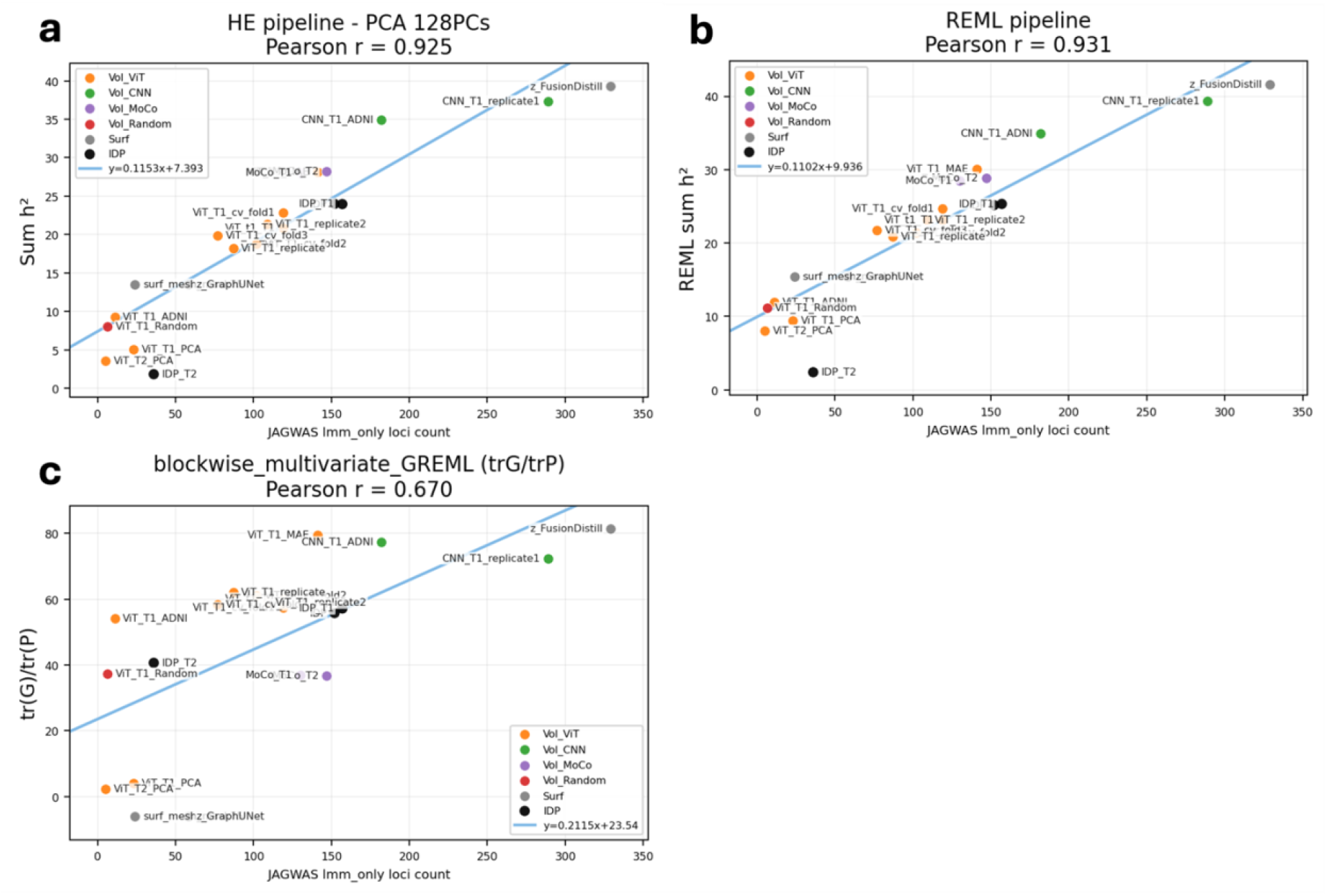
Validation of sum-h^2^ using the number of loci discovered by JAGWAS-based multivariate aggregation. (a) Base framework using a PCA cutoff of 128 PCs and HE regression for total heritability estimation. (b) Variant framework using REML for total heritability estimation while retaining PCA with a cutoff of 128 PCs. (c) Previous SOTA approach using blockwise multivariate GREML to measure total heritability with tr(G)/tr(P), compared with our measurement of total heritability as tr(P^-1^G).

As an additional validation experiment (Figure 2b), we replaced HE regression with REML for heritability estimation while keeping the remaining components of the framework unchanged. The resulting Pearson correlation was comparable to that obtained using HE regression, indicating that the proposed total heritability metric is robust to the choice of heritability estimation method.

Finally, we tested the correlation between blockwise tr(G)/tr(P) and the number of genomic loci (Figure 2c). The Pearson correlation (r = 0.670) is largely lower than that obtained from the PCA-decorrelation–HE regression framework. This is expected as tr(G)/tr(P) is not linear-invariant, so its values shift systematically across modalities (T1 → T2 → mesh) in ways that do not reflect underlying heritable signal, weakening its alignment with loci count.

We further provide theoretical explanation to this near-perfect Pearson correlation. For a multivariate association test of SNP *j* against *K* phenotypes, the test statistic asymptotically follows a non-central *χ*^2^(*K*)distribution under the alternative hypothesis (i.e., in the presence of genetic association at the locus) with non-centrality parameter 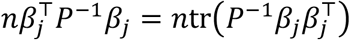, where *n* is the sample size, *P* is the phenotypic covariance matrix, and β_*j*_ is the vector of true SNP effects across the *K* phenotypes. Defining the rank-1 genetic covariance contributed by SNP *j* as 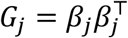,the aggregate non-centrality parameter across all causal variants is 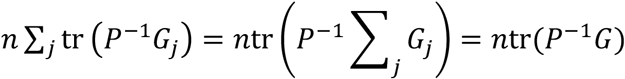, where *G* =∑ _*j*_ *G*_*j*_ denotes the total genetic covariance matrix. Thus, tr(*P*^−1^*G*)is proportional to the aggregate non-centrality parameter of an omnibus multivariate association test and quantifies the total discoverable multivariate genetic signal. Because the statistical power of multivariate GWAS methods is determined by these non-centrality parameters, tr(*P*^−1^*G*)is expected to correlate strongly with the number of loci detected under the polygenic model. Although demonstrated here using JAGWAS, this interpretation is not specific to JAGWAS and applies broadly to omnibus multivariate association frameworks, including Wald-, score-, or likelihood-ratio-based tests such as MANOVA, CCA-based methods, MOSTest, and related multivariate GWAS approaches [25-27].

The sensitivity analysis (Table 2) further shows that the Pearson correlation remains largely consistent across different sample sizes and kinship cutoffs. Specifically, correlations remained 0.92 for the JAGWAS when using: (1) kinship cutoff above 4th-degree relatives (1,470 samples), (2) kinship cutoff above 4.5th-degree relatives or 0.022 relatedness threshold (2,158 samples; default setting), and (3) kinship cutoff above 5th-degree relatives (8,126 samples). (4) kinship cutoff above 4.5th-degree relatives or 0.022 relatedness threshold with standardization preprocessing rather than demean (5) kinship cutoff above 4.5th-degree relatives or 0.022 relatedness threshold with QR decorrelation rather than PCA.

**Table 2.**
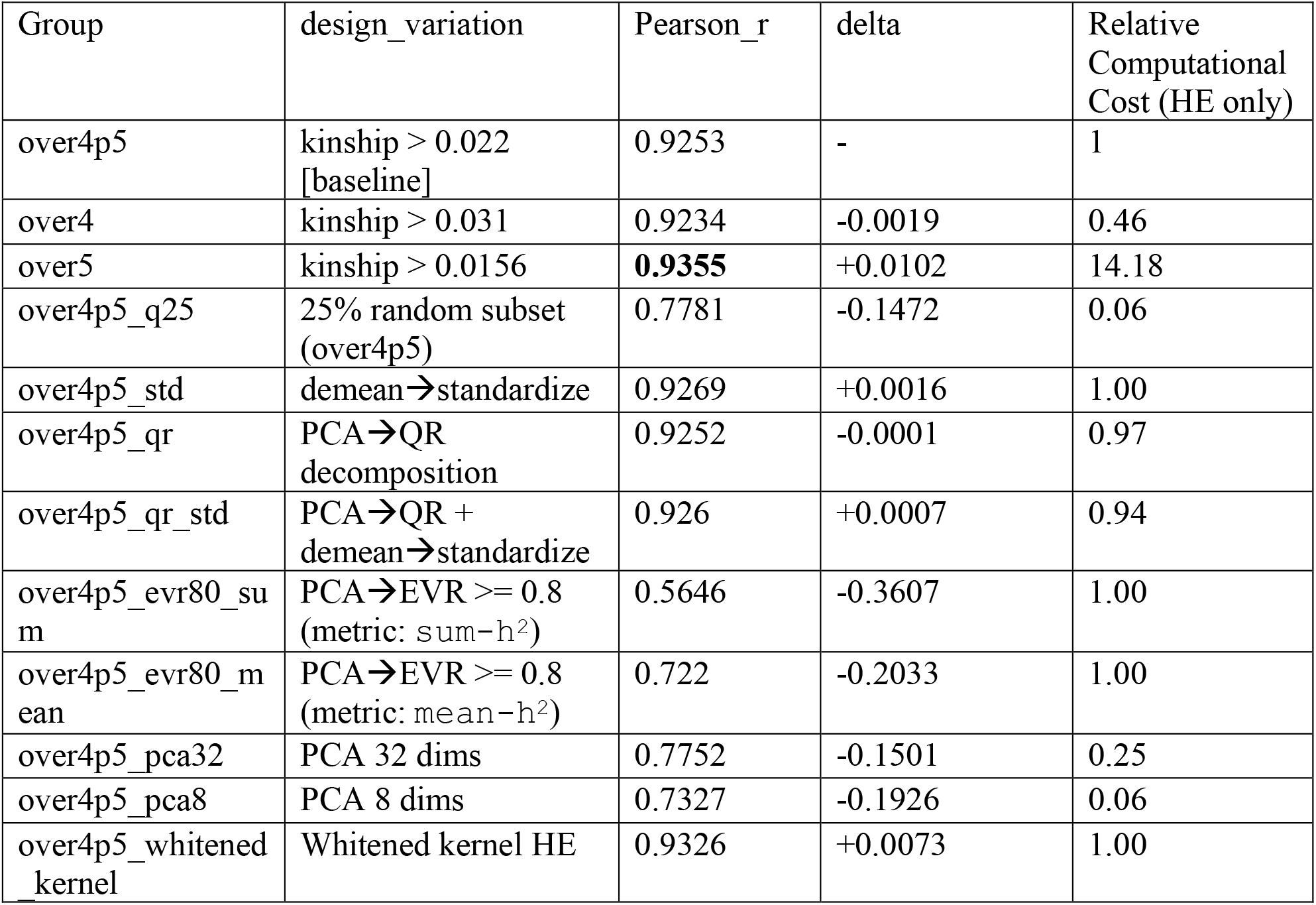
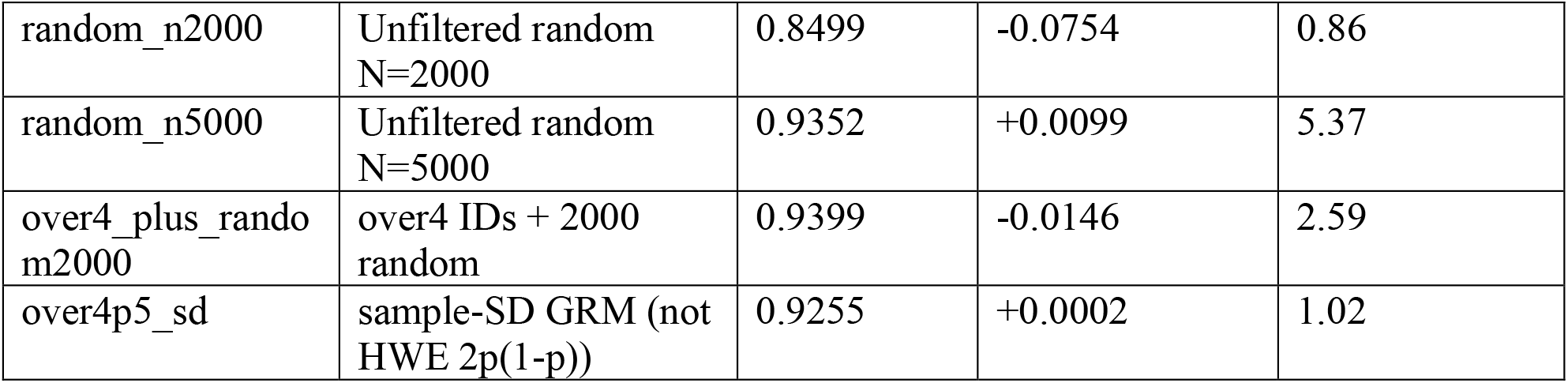
Sensitivity analysis of sum-h^2^ framework by varying different steps. Several analysis parameters were varied, including the sample-selection criterion (kinship over4th degree relatedness, over4p5, over5, or random subsampling to a matched sample size), phenotype preprocessing (demeaning or standardization), decorrelation method (PCA or QR), PCA dimensionality (fewer retained components or an explained-variance cutoff), GRM construction assumptions (with or without Hardy–Weinberg equilibrium), and a simplified one-step procedure that avoids explicit PCA through kernel whitening (see Discussion and Supplementary Note 2). Over4p5 denotes exclusion based on a kinship threshold corresponding to relatives closer than approximately 4.5th degree (>0.022); this criterion yielded 2,158 samples and was used as the default setting.

| Group | design_variation | Pearson_r | delta | Relative Computational Cost (HE only) |
| --- | --- | --- | --- | --- |
| over4p5 | kinship > 0.022 [baseline] | 0.9253 | - | 1 |
| over4 | kinship > 0.031 | 0.9234 | -0.0019 | 0.46 |
| over5 | kinship > 0.0156 | **0.9355** | +0.0102 | 14.18 |
| over4p5_q25 | 25% random subset (over4p5) | 0.7781 | -0.1472 | 0.06 |
| over4p5_std | demean → standardize | 0.9269 | +0.0016 | 1.00 |
| over4p5_qr | PCA → QR decomposition | 0.9252 | -0.0001 | 0.97 |
| over4p5_qr_std | PCA → QR + demean → standardize | 0.926 | +0.0007 | 0.94 |
| over4p5_evr80_sum | PCA → EVR >= 0.8 (metric: sum-h<sup>2</sup>) | 0.5646 | -0.3607 | 1.00 |
| over4p5_evr80_mean | PCA → EVR >= 0.8 (metric: mean-h<sup>2</sup>) | 0.722 | -0.2033 | 1.00 |
| over4p5_pca32 | PCA 32 dims | 0.7752 | -0.1501 | 0.25 |
| over4p5_pca8 | PCA 8 dims | 0.7327 | -0.1926 | 0.06 |
| over4p5_whitened_kernel | Whitened kernel HE | 0.9326 | +0.0073 | 1.00 |
| random_n2000 | Unfiltered random<br>N=2000 | 0.8499 | -0.0754 | 0.86 |
| random_n5000 | Unfiltered random<br>N=5000 | 0.9352 | +0.0099 | 5.37 |
| over4_plus_rando<br>m2000 | over4 IDs + 2000<br>random | 0.9399 | -0.0146 | 2.59 |
| over4p5_sd | sample-SD GRM (not<br>HWE 2p(1-p)) | 0.9255 | +0.0002 | 1.02 |

The comparatively lower correlation was observed in the random 25% subsampling group (q25), where the Pearson correlation was 0.78 for the JAGWAS, as well as some framework variation in the PCA process. Nevertheless, all groups still demonstrated strong positive correlations overall.

Taken together, these validation studies support the use of summed heritability following PCA-based decorrelation (sum-h^2^) as a reliable and robust metric for quantifying total heritability.

### Performance

The major advantage of our proposed total heritability estimator sum-h^2^ is its substantially improved computational efficiency in both runtime and memory usage (Figure 3). We compared our method (sum-h^2^) against an equivalent framework in which HE regression was replaced by GCTA-REML (the REML framework). We also compared it with direct computation of tr(*P*^−1^*G*), singular value decomposition (SVD) of *P*^−1^*G*, and the previously reported blockwise multivariate GREML benchmark [12], which measures total heritability as tr(*G*)/tr(*P*). Although this definition of total heritability is known to be less stable, we still included it as a comparison baseline.

**Figure 3.**
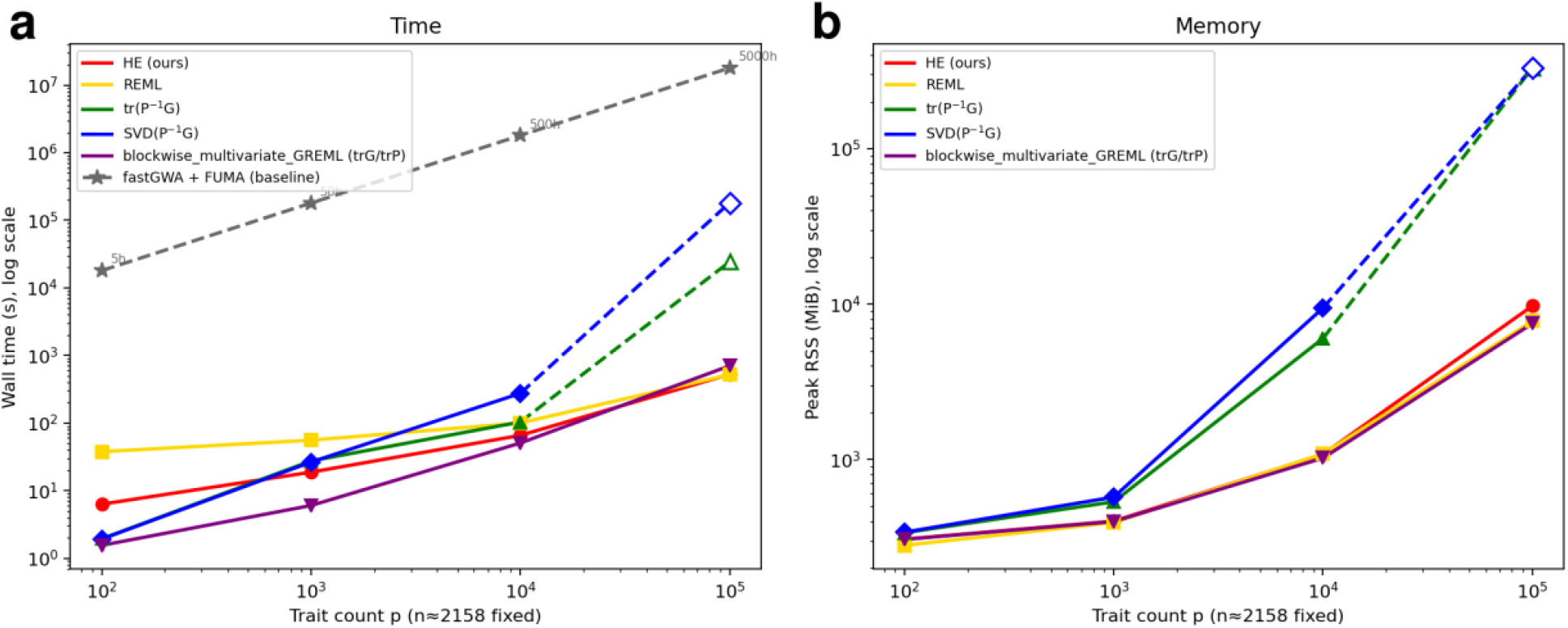
Performance of sum-h^2^ framework by different size of phenotype features (sample size fixed at 2158). (a) Time performance. (b) Memory performance.

Here is the side-by-side comparison

**Table 3.** Performance of sum-h^2^ framework by different size of phenotype features (sample size fixed at 2158).

| Analysis group | p=100 | p=1k | p=10k | p=100k time | p=100k peak RSS |
| --- | --- | --- | --- | --- | --- |
| HE framework – PCA 128PCs (ours) | 6.7s | 22.0s | 75.0s | 525.5s / 0.15h | 9.52 GiB |
| REML framework | 37.6s | 59.1s | 107.1s | 519.9s / 0.14h | 7.59 GiB |
| tr(P<sup>-1</sup>G) | 2.2s | 26.7s | 104.8s | 24,270.2s / 6.74h<br>† | 323.5 GiB † |
| SVD(P<sup>-1</sup>G) | 2.1s | 27.0s | 245.8s | 181,719.1s /<br>50.5h † | 323.5 GiB † |
| blockwise mvGREML | 2.0s | 7.1s | 60.1s | 701.6s / 0.19h | 7.32 GiB |
| fastGWA + FUMA | ~5 h † | ~50 h † | ~500 h † | ~5000 h † | - |

† estimated (see Methods)

Theoretical complexity analysis predicts that HE-PCA scales as *O*(*npk*), where *k* is the number of PCs. Because PCA fixes (k=128) regardless of the original feature dimension, its computational cost grows approximately linearly with the number of traits and remains largely insensitive to high-dimensional feature spaces. Consistent with this expectation, the HE-PCA-128PCs variation of sum-h^2^ framework (ours) is the fastest end-to-end heritability framework across all tested scales, requiring only 525.5 seconds (0.15 h) and 9.51 GiB of peak memory at (p=100,000). At large trait dimensions, the dominant computational burden arises from data loading, residualization, and PCA preprocessing, while the HE regression itself remains inexpensive due to the fixed 128-dimensional representation.

REML variation incurs a fixed per-component fitting cost that HE does not, because GCTA’s Average Information-REML algorithm must iteratively solve for variance components rather than computing them in a single pass. For each of the 128 principal components, GCTA constructs the n × n phenotypic variance matrix 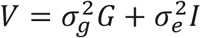and then applies the Average Information algorithm: each iteration computes the gradient and expected information matrix of the restricted log-likelihood, requiring a Cholesky factorization of V at O(n^3^) cost, and the loop repeats until convergence, typically 10-50 iterations depending on trait complexity. Running this independently across all 128 components therefore adds a block of *O*(128 × n^3^ × n_iter_)work on top of the shared residualization and PCA preprocessing. At small p this fixed overhead dominates: at p = 100, REML takes 37.6 s versus 6.7 s for HE-PCA (5.6×), the extra ∼31 s is attributable almost entirely to the 128 GCTA-REML jobs. As p grows and preprocessing time scales with the full n × p feature matrix, the iterative fitting cost becomes a shrinking fraction of the total: the ratio falls to 2.7× at p = 1k, 1.4× at p = 10k, and effectively 1.0× at p = 100k (519.9 s vs. 525.5 s), where loading and transforming the feature array accounts for nearly all wall time in both methods.

In contrast, covariance-based methods that explicitly construct the full *p* × *p* phenotypic covariance matrix exhibit substantially less favorable scaling. The trace measurement, tr(*P*^−1^*G*), requires matrix factorization and linear solves with theoretical *O*(*p*^3^)time complexity and *O*(*p*^2^)memory usage, while SVD(*P*^−1^*G*)incurs an even larger constant factor due to full eigendecomposition. Extrapolated results at *p* =100,000 suggest runtimes of 24,270 seconds (6.74 h) for tr(*P*^−1^*G*)and 181,719 seconds (50.5 h) for SVD(*P*^−1^*G*), with both methods requiring approximately 323.5 GiB of memory. These requirements render dense covariance approaches impractical for large-scale imaging-derived phenotype analyses.

Blockwise mvGREML alleviates the memory bottleneck by partitioning traits into smaller blocks, reducing the effective covariance dimension processed at any one time. Consequently, its empirical scaling remains tractable, reaching 701.6 seconds (0.19 h) at (p=100,000). However, despite this improvement, it remains slower than HE-PCA while lacking the dimensionality-reduction advantage that enables HE-PCA to maintain near-linear scaling with trait dimension.

Overall, both the theoretical complexity analysis and empirical benchmarks demonstrate that the PCA-based HE (sum-h^2^) framework provides the most favorable combination of runtime, memory efficiency, and scalability for high-dimensional heritability estimation (Table 3).

We further evaluated the impact of sample size on computational performance (Figure 4). The runtime of both HE-based sum-h^2^ frameworks is primarily driven by the sample size n. As n increases, the GRM grows quadratically in size (n x n), leading to substantially higher memory requirements and longer loading times. Because each HE regression repeatedly operates on the GRM, the cost of heritability estimation increases with sample size even when the number of traits remains unchanged.

**Figure 4.**
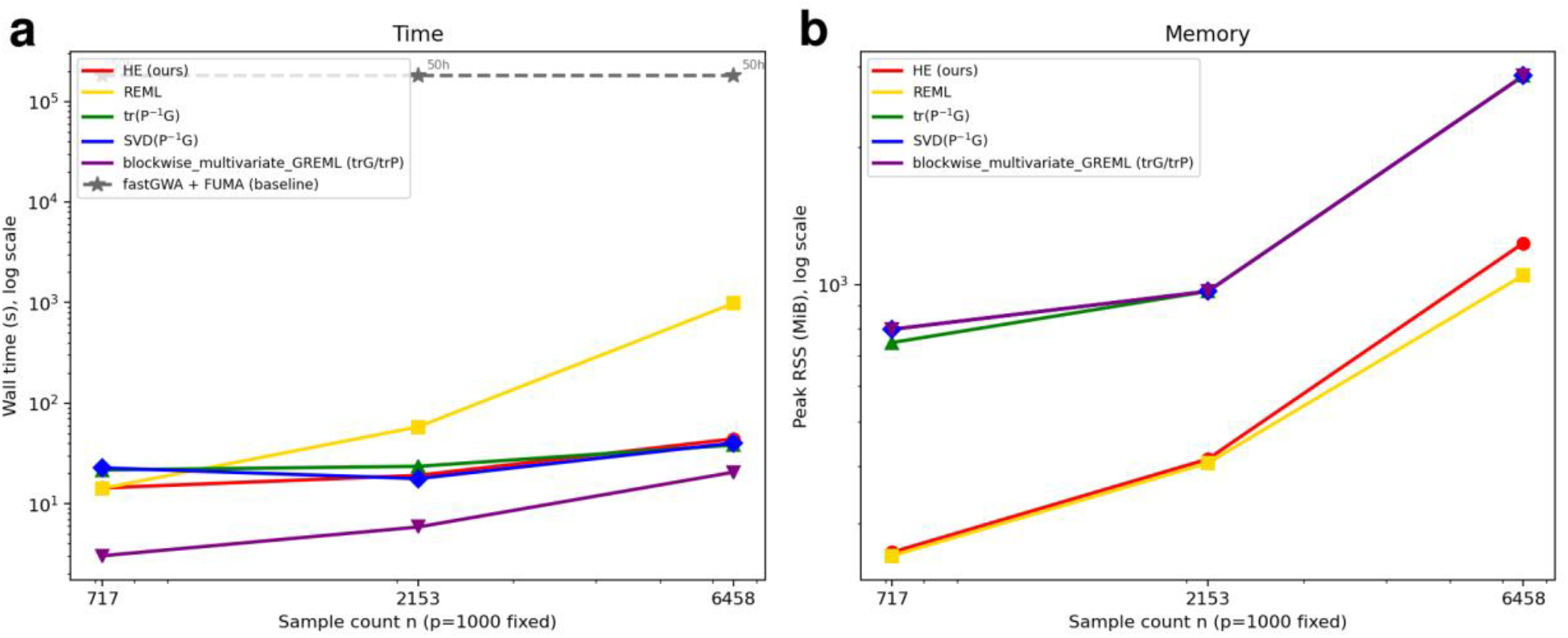
Performance of sum-h^2^ framework by different sample sizes (feature size fixed at 1000). (a) Time performance. (b) Memory performance.

### Package Support

We plan to provide full support for packaging and deploying sum-h^2^ workflow on the UKB Research Analysis Platform (RAP), enabling users to run the complete framework with minimal setup and a single-click execution once the required filtered phenotypes are ready. Due to the temporary suspension of the UKB RAP platform, full RAP integration will be released alongside the publication of this work. In the meantime, the complete source code and documentation are publicly available through GitHub. The framework repository can be accessed at: https://github.com/ZhiGroup/sum-h2

To participate in the evaluation arena, users must have authorized access to the UK Biobank genomic data under their own UK Biobank application. To ensure a fair benchmark, participants must not use the held-out test cohort during representation learning or any stage of model training. After training their representation model on the remaining eligible UK Biobank participants, users should generate representations with 128 dimensions for the held-out test cohort and evaluate them using either the RAP deployment (when available) or the open-source evaluation framework provided in the GitHub repository. For now since the RAP platform is currently unavailable, we cannot enforce the use of a shared test cohort. For a temporary solution, we ask users to apply a kinship threshold of >0.022 to exclude related individuals from all UK Biobank participants who had both brain MRI and genetic data (August 2020), resulting in a finite and deterministic test cohort of 11,963 participants after kinship filtering from 84,361 participants. Also the larger test cohort would help reduce the variability in heritability estimation, as demonstrated by the jackknife simulation (Supplementary Note 3).

For benchmarking purposes, the resulting total heritability estimates from a given model can be directly uploaded to the Overall Heritability Arena leaderboard on Hugging Face, facilitating standardized comparison across representation-learning approaches: https://huggingface.co/spaces/no1summmer/Total_heritiability_ARENA

## Discussion

This paper has three major contributions. First, we revisit the definition of total multivariate heritability in high-dimensional phenotypic systems. Most previous studies summarized global heritability using the ratio tr(*G*)/*tr*(*P*), which depends on the arbitrary scaling of traits and can change under simple linear transformations of the phenotype space. In contrast, we rediscover the quantity 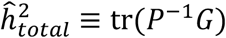, originally connected to the multivariate evolvability literature, as a more principled measurement of total additive genetic contribution 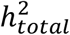. This formulation is linear invariant, accounts for phenotypic covariance structure, and can be interpreted as the multivariate analogue of narrow-sense heritability in correlated trait systems.

The second contribution of the paper is computational scalability. Direct estimation of multivariate genetic covariance matrices P and G becomes prohibitively slow and unstable in high-dimensional settings (the *P* could have phenotype highly correlated and be singular thus

*P*^−1^ is prone to error), especially for dense features such as image pixels, deep learning embeddings, or large omics profiles. To address this limitation, we introduce a “sum-h^2^” framework that combines phenotypic PCA with Haseman–Elston regression. By projecting the phenotype matrix into orthogonal principal components and estimating heritability independently for each component, the framework converts a difficult multivariate estimation problem into a set of efficient parallelizable univariate regressions. The sum of PC heritability is then exactly equivalent to tr(*P*^−1^*G*).

Finally, we propose a sum-h^2^ arena as a standardized benchmarking platform for imaging genetics and representation learning research. Because the metric is representation-invariant, different groups can directly compare phenotype representations on the same biological dataset using a common measure of total heritable signal. This enables fair evaluation between traditional imaging-derived phenotypes, handcrafted features, and modern deep learning embeddings. We will have the supporting package ready on the UKB-RAP platform as well as the leaderboard/arena ready on the hugging face.

Several design choices in the proposed framework require further clarification. First, we construct a subsample of approximately 2,000 individuals with a kinship threshold of 0.022. Although the default UK Biobank framework often uses 0.044 to derive a sparse GRM, our goal is to obtain a denser GRM with sufficient off-diagonal signal, which motivates a more permissive cutoff. At the same time, we retain enough samples for stable estimation, making 0.022 a balanced choice. We also tested alternative thresholds in simulation studies and found that results remain highly consistent and closely approximate the full-sample total heritability. For GRM construction, we use GCTA under the HWE assumption, which empirically produces the most accurate dense GRM compared with direct computation approaches, particularly when evaluated against IDP-based benchmarks.

Second, the framework includes options for phenotype preprocessing and covariate adjustment. By default, we use mean-centering (demeaning) of phenotypes, but users may choose standardization when combining features with heterogeneous scales to improve numerical stability. Covariate inclusion is also configurable; we currently include T1-specific covariates to maintain consistency with prior work, although these can be modified depending on the modality or research goal. Finally, the current total heritability metric ranges from 0 to 128 due to summation across 128 PCs, and we are considering normalizing by the number of components to rescale the measure into the conventional [0,1] range for interpretability.

Furthermore, there could be an optimization to further improve algorithmic efficiency by condensing the two-step framework (PCA followed by HE regression) into a direct, one-step HE regression (Supplementary Note 2). This is achieved by evaluating the *n* × *n* phenotypic similarity matrix *S* =*YP*^−1^*Y*^⊤^, where *Y* is the *n* × *k* phenotype matrix and *P* is the *k* × *k* phenotypic covariance matrix. A single HE regression of the off-diagonal elements of *S* onto the *n* × *n* genetic relatedness matrix (GRM) directly yields tr(*P*^−1^*G*)as the slope (detailed proof provided in Supplementary Note 2). For moderate feature spaces (*k* =100), this one-step approach is approximately four times faster than the two-step PCA framework. However, this computational advantage diminishes as the feature space grows; for large *k*, empirical covariance estimation becomes the primary temporal bottleneck alongside increased memory demands for storing *P*. Moreover, the direct method requires the explicit computation of *P*^−1^, which becomes numerically unstable if *P* is ill-conditioned (e.g., as *k* approaches *n*). The PCA-based approach naturally circumvents this vulnerability by allowing the truncation of principal components with near-zero eigenvalues, avoiding the inversion of near-singular matrices entirely. Ultimately, this direct one-step formulation provides a mathematically elegant alternative that is highly amenable to future GPU acceleration to further optimize the underlying matrix multiplications.

## Methods

### UK Biobank

The UK Biobank [28] is a nationwide population resource encompassing approximately 500,000 participants recruited across the United Kingdom, with phenotypic, imaging, and genetic data collected between 2006 and 2010. The study operates under established ethical approval, with governance and oversight details available at https://www.ukbiobank.ac.uk/learn-more-about-uk-biobank/governance/ethics-advisory-committee. In total, 29,115 brain MRI scans were included in this study. Model development used 4,597 scans for training and 1,533 for validation. Learned representations were subsequently extracted from the remaining 22,985 individuals, corresponding to the same “discovery cohort” used in prior brain imaging GWAS analyses [29, 30].

For genetic analyses, imputed genotype data from the UK Biobank [28] underwent standard quality control procedures. Variants were retained if they met thresholds of minor allele frequency > 0.0001, resulting in 8,925,988 single-nucleotide polymorphisms in the discovery cohort.

### Alzheimer’s Disease Neuroimaging Initiative (ADNI) [31]

Data used in the preparation of this article were obtained from the Alzheimer’s Disease Neuroimaging Initiative (ADNI; including ADNI-1, ADNI-2, ADNI-3, and ADNI-GO; downloaded in February 2025) database (adni.loni.usc.edu). The ADNI was launched in 2003 as a public-private partnership, led by Principal Investigator Michael W. Weiner, MD. The primary goal of ADNI has been to test whether serial magnetic resonance imaging (MRI), positron emission tomography (PET), other biological markers, and clinical and neuropsychological assessment can be combined to measure the progression of mild cognitive impairment (MCI) and early Alzheimer’s disease (AD). A total of 2,594 MRI scans from 1,524 subjects were included in this study.

### Model Input Datasets and Preprocessing

UK Biobank MRIs were downloaded on October 15, 2021. UK Biobank participants had birth year between 1934 to 1971 with female to male ratio of 0.52. UK Biobank has provided a bias-field-corrected version of the brain-extracted T1-weighted (T1) captured mainly using standard Siemens Skyra 3T running VD13A SP4 (as of October 2015), with a standard Siemens 32-channel RF receive head coil.

Resolution of T1 is 1 × 1 × 1 mm (https://biobank.ctsu.ox.ac.uk/crystal/crystal/docs/brain_mri.pdf).

To promote generalizability and minimize manual feature engineering, we adopted the standard preprocessing pipeline developed by the UK Biobank MRI team. Brain MRI preprocessing was performed primarily using FSL (https://www.fmrib.ox.ac.uk/ukbiobank/), and included defacing, brain extraction with BET, linear and non-linear registration to standard space using FLIRT and FNIRT, and bias-field correction using FAST. Bias-field-corrected, brain-extracted T1-weighted images were retained for analysis. All images were subsequently linearly registered to MNI152 space using affine transformation with 12 degrees of freedom via UK Biobank-provided precomputed matrices in FLIRT. Linear registration was chosen to normalize head size and achieve cross-subject alignment while preserving subject-specific structural deformations, in contrast to non-linear registration. The resulting linearly registered, defaced, bias-field-corrected images were used in all downstream analyses. Image intensities were normalized on a per-scan basis using Z-score normalization. Following affine registration, T1 weighted volumes had dimensions of 182 × 218 × 182 voxels and were zero-padded to 182 × 224 × 182 to enable partitioning into equal-sized patches.

To construct a high-quality deep learning dataset, we leveraged UK Biobank’s precomputed MRI quality metrics: the inverted contrast-to-noise ratio (Data Field 25735) and the discrepancy between T2 FLAIR and T1 brain images (Data Field 25736). Images with values below the 95th percentile for both metrics-corresponding to higher-quality scans-were retained. For participants with multiple visits, only the first scan was kept to ensure consistency across the dataset. This filtering resulted in a cohort of 6,130 scans from individuals of diverse ethnic backgrounds. The dataset was randomly split into a training set of 4,597 images (75%) and a validation set of 1,533 images (25%). The validation set was reserved exclusively for hyperparameter tuning and model checkpointing during training.

For the ADNI data, we downloaded 2,594 T1-weighted structural MRI scans acquired on 3T scanners in NIfTI format, each with vendor-provided on-scanner intensity non-uniformity correction. All images were processed using an in-house preprocessing pipeline to ensure consistency across sites and acquisition protocols. First, skull stripping was performed using DeepBET (threshold = 0.5) to remove non-brain tissue while preserving cortical boundaries [32]. Next, images were spatially normalized to the MNI152 standard space via affine registration using ANTs [33], enabling cross-subject anatomical alignment. Finally, N4 bias field correction (implemented in ANTs) was applied to further reduce residual intensity inhomogeneity. This standardized preprocessing pipeline facilitates robust downstream feature extraction and cross-cohort comparability. The final T1 weighted volumes had dimensions of 182 × 218 × 182 voxels and resolution of 1 × 1 × 1 mm and were zero-padded to 182 × 224 × 182 to enable partitioning into equal-sized patches. The dataset was randomly split into a training set of 1,945 images (75%) and a validation set of 649 images (25%).

### Model training

We adopt CNN [20]and ViT [34] autoencoder framework from our previous publication. Details could be found in our previous paper.

Briefly, The CNN architecture is an autoencoder with CNN block of 3 × 3 × 3 kernel and 2 × 2 × 2 kernel for max pooling. The model consist of 5 encoding CNN blocks, an average pooling layer with dimension 128 and 5 decoding CNN blocks. The model is trained for 100 epochs and converge around 20 epochs.

The ViT-AE architecture consisted of encoders, an average pooling layer and decoders. The raw images are partitioned into patches and embedded into 128 dimensions, fed into the vision transformer along with its position embeddings. The combined token embeddings passed through 12 encoding transformer layers, the average pooling layer, 12 decoding transformer layers and were finally decoded by a linear projection to reconstruct the input patches. MSE loss was computed between predicted patches and original patches within the batch mask. The learning schedule is a CosineAnnealingWarmRestarts learning rate scheduler with T_0=10, T_mult=2, eta_min=1×10-6 and AdamW optimizer with an initial learning rate of 0.001. The ViT-AE model was trained for 300 epochs on the training set and validated on a separate validation set on 4 Nvidia H100 GPUs.

### Model embedding extraction

We froze the CNN/ViT encoder and average pooling layer and remove the decoder layer, then applied the model to T1 weighted MRIs of 22,985 held-out individuals (discovery cohort) to get the representations of size 22,985 samples × 128 dimensions from the average pooling layer.

### Genetic analyses

We implemented a comprehensive quality-control pipeline on the downloaded genetic data, following our previous work [20], resulting in 8,469,833 single-nucleotide polymorphisms (SNPs) across 22,985 individuals of white British ancestry. Briefly, duplicated variants across the 22 autosomal chromosomes were removed, and individuals whose genetically inferred sex did not match their self-reported sex were excluded. Additional filtering criteria removed variants with a minor allele frequency (MAF) below 0.01% or a genotyping missing rate exceeding 5%.

### GWAS

GWAS were conducted on 22,867 individuals for 128 extracted UDIPs. Associations between SNPs and UDIPs were tested using linear mixed models implemented in FastGWA [22] within GCTA (v1.94.1), with a MAF threshold of 0.01. Covariates included age (Field ID: 21003), age^2^, sex (Field ID: 31), sex × age, sex × age^2^, the first 10 genetic principal components (Field ID: 22009), intracranial volume (Field ID: 25000), inverted contrast-to-noise ratio (Field ID: 25735), head positioning in the scanner (Field IDs: 25756-25758), scanner table position (Field ID: 25759), assessment center location (Field ID: 54), and date of assessment (Field ID: 53). For participants with multiple visits, only the first scan was retained. A Bonferroni-corrected significance threshold of 5 × 10^−8^/128 was applied, and summary statistics were compiled with the JAGWAS for each SNP.

### Annotation of genomic loci

Genomic loci were annotated using FUMA [4]. Lead SNPs were first identified based on linkage disequilibrium (r^2^ ≤ 0.1) and physical proximity (<250 kb), and assigned to non-overlapping genomic loci. Within each locus, the SNP with the lowest p-value (the top lead SNP) was used to represent the locus.

### JAGWAS

Joint Analysis of Multiple Phenotypes (JAGWAS) is a new tool that can efficiently calculate multivariate association statistics using single-phenotype summary statistics for hundreds of phenotypes [23]. Let *z*_*ij*_ be the test statistic (z-score) obtained from the single-phenotype GWAS between the ith phenotype and jth SNP. Then let ***Z***_***j***_ = [*Z*_1*j*_,…, *Z*_*kj*_] be the vector of z-scores across the *k* phenotypes on the jth SNP. The JAGWAS method is based on a multivariate test statistic 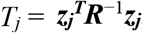, where ***R*** represents the phenotypic correlation matrix. Under the null hypothesis, this statistic asymptotically follows a chi-square distribution with k degrees of freedom. Therefore JAGWAS performs a chi-square test using the test statistic 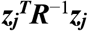, where ***R*** is estimated by the k-by-k observed correlation matrix of scaled residuals. The scaled residuals are residuals obtained through a linear mixed model, divided by the residual variance estimate.

### Validation

To evaluate the proposed total heritability metric, calculated as the sum of feature-level heritability, we assessed its association with the gold-standard indicator of genetic signal: the number of genome-wide significant loci identified in the corresponding GWAS analyses described above.

We evaluated a variant of our framework that replaces HE regression with REML for calculating hertiability. We also validated an alternative definition of total heritability, tr(G)/tr(P), and assessed its association with genetic discovery. In addition, we conducted ablation studies to evaluate the robustness of the framework under varying sample kinship structures and sample sizes.

### Performance

We evaluated the computational efficiency of our method in terms of runtime and memory usage across varying sample sizes (719, 2,158, and 6,474; 717, 2,153 and 6,458 after covariate sample ID filtering) and feature dimensions (100, 1,000, 10,000, and 100,000) using randomly generated data. The evaluation is performed on the CPU Intel Xeon Gold 6442Y, a 4th Gen Xeon Scalable (Sapphire Rapids) with 8 parallel subprocesses.

For the dense baseline methods, tr(P^−1^G) and SVD(P^−1^G), the results at 100,000 features were estimated rather than directly measured. Specifically, runtimes at 100, 1,000, and 10,000 features were transformed into log_10_–log_10_ space (log trait count versus log wall time), and a quadratic polynomial was fitted to these three observations. The fitted model was then evaluated at log_10_(100,000) to extrapolate the expected runtime. For tr(P^−1^G), a lower bound based on the theoretical O(p^3^) dense Cholesky decomposition cost was additionally imposed, and the final estimate was taken as the maximum of the polynomial prediction and this analytical dense-kernel estimate. The SVD(P^−1^G) estimate used the same lower bound scaled by the empirical runtime ratio between SVD and trace computation (approximately 7.5×) observed at smaller feature dimensions.

Memory consumption at 100,000 features for both dense methods was estimated using the corresponding dense O(p^2^) matrix storage requirement, yielding approximately 323 GB. This reflects the memory needed to construct and factorize a full 100,000 × 100,000 covariance matrix stored in double precision.

For the GWAS analysis, fastGWA and FUMA were benchmarked using ∼20,000 samples, a cohort size that is representative of a typical GWAS with sufficient statistical power to detect biologically meaningful associations. The fastGWA analysis were performed with 8 parallel jobs, same as the HE-PCA analysis. Since time for running FUMA is not stable and requires manual upload to the website, we give an empirical estimate of the total time with fastGWA and FUMA together.

The reported results for HE-PCA, HE-QR, REML, and blockwise mvGREML at 100,000 features were obtained through direct execution. These methods avoid constructing the full feature-by-feature covariance matrix and therefore remain computationally feasible at this scale. Consequently, the runtime and memory measurements reported for these approaches at 100,000 features represent actual end-to-end benchmark results rather than extrapolated estimates.

## Data Availability

The previous published CNN and ViT model codes as well as checkpoints are publicly accessible at Github at https://github.com/ZhiGroup/DeepENDO, https://github.com/ZhiGroup/UDIP-ViT.

The UDIPs and GWAS summary statistics are available upon reasonable request after removing sensitive UKbiobank patient information. Genomic loci annotation used data from FUMA (https://fuma.ctglab.nl/). Individual data from UKBB can be requested with proper registration at https://www.ukbiobank.ac.uk/. All unrestricted data supporting the findings are also available from the corresponding author upon request.

## Acknowledgements

This work was supported by grants from the National Institute on Aging U01AG070112 and R01AG081398.

## Ethics declarations

### Ethical Approval

Our analysis was approved by the UTHealth Houston committee for the protection of human subjects under No. HSC-SBMI-20-1323. UKBB has secured informed consent from the participants in the use of their data for approved research projects. UKBB data was accessed via approved project 24247.

